# MO25 binds CBL-interacting protein kinases associated with ribonucleoprotein condensates and regulates meiotic exit

**DOI:** 10.64898/2026.08.19.745790

**Authors:** Anna Vargova, Jana Faturova, Albert Cairo, Kristyna Jankujova, Jana Pecinkova, Pavlina Mikulkova, Claudio Capitao, Karel Riha

## Abstract

Meiotic (M)-bodies are multiphasic ribonucleoprotein (RNP) condensates composed of a P-body core surrounded by a stress granule-like shell that promote meiotic exit through transient translational repression. This process depends on the phosphoserine-binding protein SMG7, which recruits the meiotic regulator TDM1 to M-bodies during meiosis II. Here, we identify the evolutionarily conserved scaffold protein MO25 as a regulator of SMG7 and TDM1 partitioning into M-bodies in *Arabidopsis thaliana*. Disruption of MO25A1 enhances the accumulation of SMG7 and TDM1 in M-bodies and increases the reduced fertility in the hypomorphic *smg7-6* mutant, which exhibits impaired M-body association. In fungi and animals, MO25 proteins act as allosteric activators of STE20-family kinases. Interaction screening revealed that, whereas Arabidopsis MO25B proteins interact with STE20-family MAP4K kinases, MO25A paralogues have evolved specificity toward a subset of CBL-interacting protein kinases (CIPKs). Notably, the MO25A-interacting CIPKs localize to diverse nuclear and cytoplasmic RNP condensates. Among them, CIPK6 is required for fertility and pollen development, and disruption of its MO25-binding domain enhances SMG7 condensation. Together, our findings identify a previously unrecognized MO25A–CIPK interaction module that regulates M-body organization and may more broadly contribute to the regulation of RNP condensates.

## Introduction

Ribonucleoprotein (RNP) condensates are microscale subcellular compartments that function as hubs for RNA storage and processing. They are highly dynamic structures that exhibit properties of liquid– liquid phase-separated condensates, enabling them to rapidly alter their composition and biochemical properties in response to changes in cellular physiology (Ripin and Parker, 2023; Maity and Moschou, 2026). RNP condensates have emerged as key regulators of processes that require extensive remodeling of gene expression, including developmental transitions and cellular stress responses.

Among the best-characterized cytoplasmic RNP condensates in plants are processing bodies (P-bodies) and stress granules (SGs), which play central roles in translational regulation, RNA storage, and mRNA decay (Chantarachot and Bailey-Serres, 2018; Solis-Miranda et al., 2023; Solis-Miranda et al., 2026). SGs form in response to stress-induced translational arrest, when mRNAs released from disassembling polysomes condense together with cytoplasmic RNA-binding proteins. Once the stress subsides and translation resumes, SGs disassemble. In contrast to SGs, P-bodies are constitutively present in the cytoplasm, although their abundance and size are dynamically regulated by cellular conditions. P-bodies are enriched in translationally repressed mRNAs and proteins involved in mRNA decay, including decapping enzymes, exonucleases, and components of the RNA-induced silencing machinery. Under stress conditions, SGs and P-bodies are closely interconnected. They frequently dock or transiently associate, suggesting coordinated regulation of mRNA storage, translation, and decay (Chantarachot and Bailey-Serres, 2018; Ripin and Parker, 2023).

SGs are primarily involved in stress adaptation and recovery, helping plants survive abrupt environmental changes and rapidly resume growth once stress ends (Wang et al., 2024; Xie et al., 2024; Geng et al., 2025; Melicher et al., 2026; Solis-Miranda et al., 2026). P-bodies function more broadly, regulating developmental transitions, hormonal responses, virus infection and immune signaling (Jang et al., 2019; Yu et al., 2019; Chantarachot et al., 2020; Cairo et al., 2022; Hoffmann et al., 2022; Liu et al., 2023; Liu et al., 2024). The remarkable plasticity of SGs and P-bodies stems from the weak multivalent interactions that hold their components together. Reversible phosphorylation can modulate these interaction networks, enabling signaling pathways to rapidly reshape condensate behavior in response to developmental and environmental cues (Wippich et al., 2013; Protter and Parker, 2016; Sfakianos et al., 2018; Legoux et al., 2025). However, kinase-driven regulation of stress granules and P-bodies in plants is only beginning to be understood. Key examples include the MAP kinases MPK3 and MPK6, which have been implicated in P-body disassembly and SG homeostasis during pathogen-induced signaling (Yu et al., 2019; Tabassum et al., 2020), as well as sucrose- non-fermenting 1-related kinases 1 and 2 (SnRK1/2), whose activity is regulated through condensation within SGs and P-bodies (Gutierrez-Beltran et al., 2021; Yuan and Zhao, 2025).

Phosphorylation has also been implicated in the regulation of meiotic (M)-bodies, specialized multiphasic RNP condensates comprising a P-body core, an SG-like shell, and a CDM1-rich phase formed by the RNA-binding protein CDM1 (Saddala et al., 2025; Cairo et al., 2026). M-bodies are important regulators of meiosis, and their composition and functional properties change dynamically during meiotic progression. At the end of meiosis, M-bodies recruit TDM1, a tetratricopeptide repeat protein that sequesters the translation initiation factor eIFiso4G2, thereby temporally inhibiting translation. This process is crucial for the termination of meiosis and the transition to post-meiotic pollen development (Cairo et al., 2022). Premature recruitment of TDM1 to M-bodies is inhibited by the meiotic cyclin TAM, which, together with cyclin dependent kinase CDKA;1, phosphorylates TDM1 (Cifuentes et al., 2016; Schindfessel et al., 2026). During heat stress, TAM itself is sequestered into M-bodies, preventing premature meiotic termination and the formation of diploid pollen (De Jaeger-Braet et al., 2025; Schindfessel et al., 2026).

The importance of protein phosphorylation in M-body regulation is further highlighted by the central role of SMG7, an evolutionarily conserved P-body component that recognizes phosphorylated proteins through its 14-3-3-like domain (Fukuhara et al., 2005). SMG7 supports M-body function through at least two mechanisms. First, it mediates the recruitment of TDM1, thereby promoting meiotic exit (Cairo et al., 2022). Second, it maintains M-body size homeostasis by recruiting eIF4A helicases, RNA chaperones that remodel RNA–RNA and RNA–protein interactions within RNP condensates (Tauber et al., 2020; Cairo et al., 2026). Importantly, the interaction of SMG7 with both TDM1 and eIF4A helicases is thought to be regulated by phosphorylation. Together, these findings suggest that phosphorylation-dependent interactions constitute one of the key mechanisms governing M-body composition and point to the existence of kinases that dynamically modulate M-body function.

In this study, we identified MO25 as a SG component and a regulator of M-bodies. MO25 is an evolutionary conserved scaffold protein best known for its role in the LKB1-STRAD-MO25 kinase complex in animals (Boudeau et al., 2003). By binding the STRAD pseudokinase, MO25 stabilizes the heterotrimeric complex and allosterically activates the LKB1 kinase (Zeqiraj et al., 2009). Beyond the LKB1-STRAD complex, MO25 interacts with and activates several other members of the STE20 family, including SPAK/OSR and MST3/MST4/YSK1 (Filippi et al., 2011). MO25 proteins are also present in budding and fission yeasts, where they associate with kinases orthologous to STE20 family members (Nelson et al., 2003; Kanai et al., 2005). Plant MO25 homologs have also been identified, and loss of MO25A1 function in rice results in embryonic lethality (Zermiani et al., 2015; Bizotto et al., 2018; Ta et al., 2023). However, the kinase partners of MO25 in plants have yet to be identified.

Here, we identified *MO25A1* in a suppressor screen for mutations that restore fertility in *Arabidopsis smg7-6* mutants. We found that MO25A1 localizes to SGs and M-bodies under heat stress and negatively regulates the partitioning of SMG7 and TDM1into M-bodies. We further show that MO25A1 interacts with a subset of CBL-interacting protein kinases (CIPKs), including CIPK6, which associates with P-bodies and is required for pollen development. These findings uncover an unexpected role for MO25A1 in the regulation of cytoplasmic RNP granules and suggest a functional connection between CIPK signaling and RNP condensate dynamics.

## Results

### Mutation in MO25A1 improves fertility of Arabidopsis smg7-6 mutans

SMG7 associates with the P-body core of M-bodies and is essential for the completion of meiosis by recruiting TDM1 through its N-terminal 14-3-3-like domain (Riehs et al., 2008; Cairo et al., 2022; Cairo et al., 2026). Partitioning of SMG7 into M-bodies is facilitated by its intrinsically disordered C-terminal region, which contains a prion-like domain. Truncation of the C-terminal region in the *smg7- 6* allele reduces the localization of SMG7 to M-bodies and impairs TDM1 recruitment (Cairo et al., 2022). Consequently, *smg7-6* mutants phenocopy TDM1 loss of function, resulting in aberrant cycles of meiotic division and the formation of polyads (Capitao et al., 2021). This substantially reduces pollen count and fertility in *smg7-6* mutants.

To identify genes involved in the SMG7-TDM1 pathway, we performed a suppressor screen for mutations that improve *smg7-6* fertility (Capitao et al., 2021). One of the identified lines, designated *EMS225*, exhibited increased seed production, manifested by longer siliques and a higher pollen yield compared with *smg7-6* (Figure 1a-c). However, *EMS225* only partially rescued the *smg7-6* phenotype and did not restore the fertility to wild type levels.

**Figure 1.**
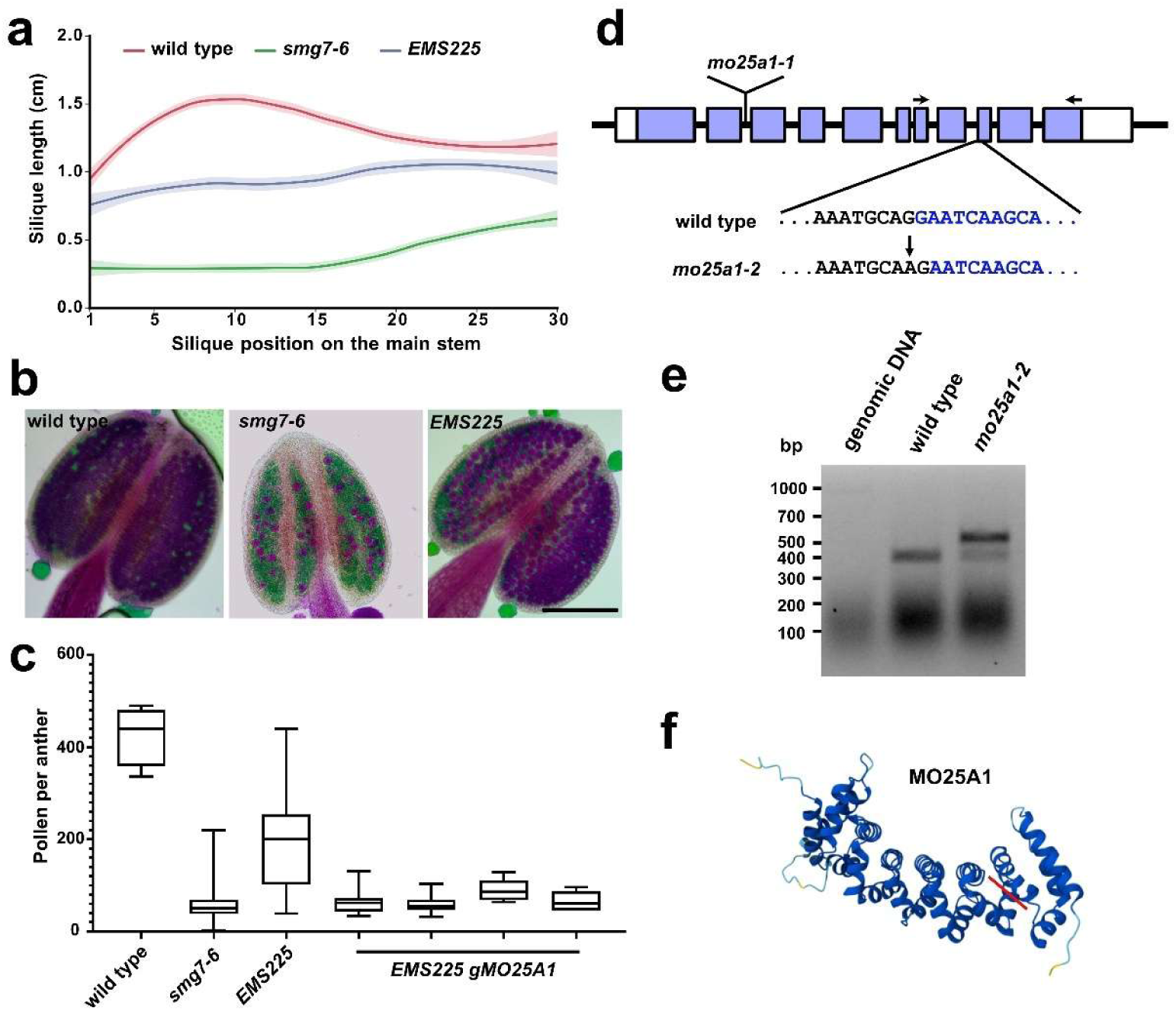
Identification of MO25A1 in a suppressor screen for enhanced fertility of *smg7-6*. a) Quantification of silique length along the main inflorescence bolt in wild type (n = 15), *smg7-6* (n = 18), and *EMS225* (n = 20). Data trends were fitted using locally weighted scatterplot smoothing (LOESS, colored lines) and shaded area indicate 95% confidence intervals. b) Anthers of indicated genotypes after Alexander staining. Scale bar = 0.1 mm. c) Box-and-whisker plot showing the number of viable pollen per anther (wild type n = 15, *smg7-6* n = 154, EMS225 n = 15, EMS225 gMO25A1 (n = 5 each line). d) Schematic representation of the *MO25A1* gene (At4g17270), showing the mutation identified in *mo25a1-2* and the position of the T-DNA insertion in *mo25a1-1* allele. Exons and are shown in blue. Arrows indicate the positions of primers used for RT-PCR analysis. e) Agarose gel electrophoresis of RT-PCR derived from *MO25A1* transcripts in wild type and *mo25a1-2* plants. f) AlphaFold-predicted structure of the MO25A1 protein. The red line indicates the position of the C- terminal truncation caused by the *mo25a1-2* mutation.

The suppressor phenotype behaved as a recessive trait and association mapping combined with whole-genome sequencing revealed a mutation in the At4g17270 locus, which encodes MO25A1 protein (Figure S1, Table S1). We designated this allele *mo25a1-2*. Transformation of *EMS225* plants with a genomic fragment corresponding to the At4g17270 locus reduced pollen production to levels similar to those observed in *smg7-6* plants (Figure 1c), demonstrating that the mutation is causal.

The *mo25a1-2* allele carries a G-to-A transition in the -1 position of the 3′ splice acceptor site of the eighth intron (Figure 1d). RT-PCR analysis followed by sequencing revealed aberrant splicing of the eighth intron, resulting in two splice variants (Figure 1e). The predominant splice variant retained the intron, whereas the second variant utilized an alternative splice site shifted by a single nucleotide.

Both splice variants introduced a frameshift and a premature termination codon shortly downstream of exon 8 and are therefore predicted to encode truncated proteins lacking approximately the C-terminal quarter of the protein (Figure 1f).

### MO25A1 and MO25A2 exhibit partial functional redundancy

To further characterize MO25A1, we obtained a T-DNA insertion line, designated *mo25a1-1*, in which the insertion disrupts the gene after the second intron and likely results in a complete loss-of-function allele (Figure 1d). Homozygous *mo25a1-1* mutants appeared phenotypically normal and were fully fertile (Figure 2). Land plants contain two distinct clades of MO25 genes, MO25A and MO25B (Bizotto et al., 2018). The *Arabidopsis thaliana* genome contains four *MO25* genes, of which At5g18940 belongs to the MO25B clade, whereas At4g17270 (*MO25A1*), At5g47540 (*MO25A2*) and At2g03410 (*MO25A3*) belong to the MO25A clade. *MO25A1* and *MO25A2* arose from a relatively recent duplication event within the *Brassicaceae* family (Bizotto et al., 2018), suggesting potential functional redundancy.

**Figure 2.**
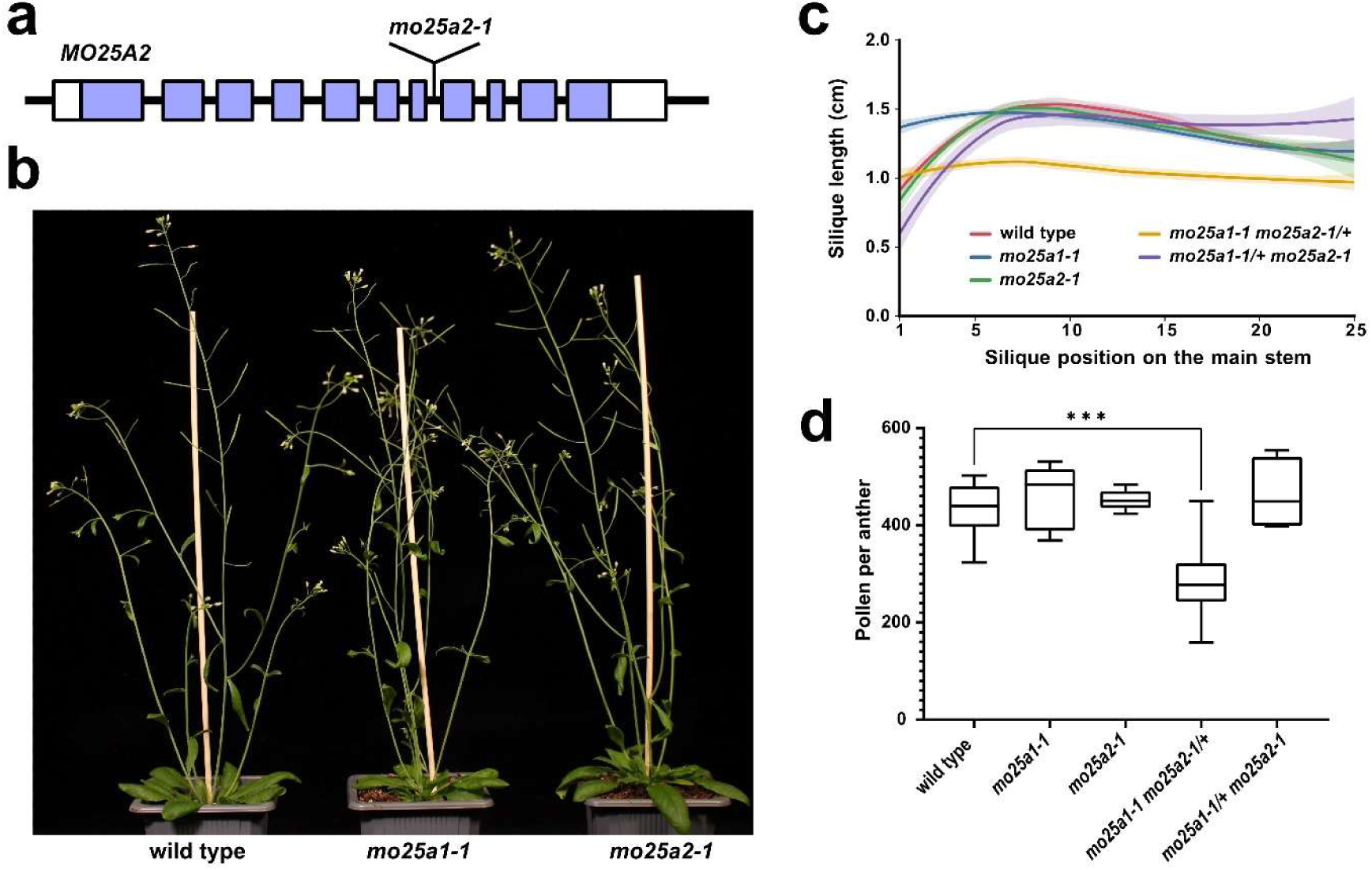
Characterization of *mo25a2-1* mutants. a) Schematic representation of the *MO25A2* gene (At5g47540), showing the position of the T-DNA insertion in the *mo25a2-1* allele. Exons are shown in blue. b) Representative phenotypes of 6-week-old wild type, *mo25a1-1,* and *mo25a2-1* plants. c) Quantification of silique length along the main inflorescence bolt in plants of the indicated genotypes; wild type (n = 15), *mo25a1-1* (n = 9), *mo25a2-1* (n = 6), *mo25a1-1 mo25a2-1/+* (n = 5), and *mo25a1-1/+ mo25a2-1* (n = 5). Data trends were fitted using locally weighted scatterplot smoothing (LOESS, colored lines) and shaded area indicate 95% confidence intervals. d) Box-and- whisker plot showing the number of viable pollen per anther (*** P ˂ 0.001, Kruskal Wallis with Dunn posthoc test, wild type (n = 15), *mo25a1-1* (n = 6), *mo25a2-1* (n = 6), *mo25a1-1 mo25a2-1/+* (n = 23), and *mo25a1-1/+ mo25a2-1* (n = 6).

To address this question, we characterized a T-DNA disruption allele of *MO25A2* (*mo25a2-1*, Figure 2a) and attempted to generate plants deficient in both MO25A1 and MO25A2. Similar to *mo25a1-1*, *mo25a2-1* plants were phenotypically normal and fully fertile (Figure 2b). However, we were unable to obtain *mo25a1-1 mo25a2-1* double mutants. Furthermore, *mo25a1-1 mo25a2-1/+* plants exhibited reduced fertility and decreased pollen formation (Figure 2c,d), although they did not display obvious vegetative growth defects. In contrast, *mo25a1-1/+ mo25a2-1* plants were fully fertile, suggesting that the contribution of *MO25A1* to fertility is more pronounced than that of *MO25A2*. To determine whether the failure to obtain double mutants was caused by defects in gametophyte development, we performed reciprocal crosses between *mo25a1-1 mo25a2-1/+* and wild type and analyzed the transmission of the *mo25a2-1* allele. Transmission through the female germline did not differ significantly from the expected 1:1 ratio (38:37; χ² = 0.013, P = 0.908). In contrast, transmission through the male germline was significantly reduced, but not fully abolished (27:51, χ² = 7.385, P = 0.007). These data suggest that *MO25A1* and *MO25A2* function redundantly and that the simultaneous loss of both genes impairs male gametophyte function and results in embryonic lethality. These data are consistent with observations in rice, where *osmo25a1* mutants also arrest during embryonic development (Ta et al., 2023).

Next, we performed protein localization studies. Transient expression of MO25A1:YFP and MO25A2:YFP in mesophyll protoplasts resulted in very similar localization patterns, with signals detected in both the cytoplasm and nucleus (Figure 3a). However, a more nuanced pattern emerged when analyzing roots of plants stably expressing genomic clones C-terminally tagged with RFP under the control of their native promoters. While MO25A2:RFP localized exclusively to the cell periphery, MO25A1:RFP additionally exhibited an intracellular signal concentrated in putative nuclei (Figure 3b). Both proteins co-localized with the plasma membrane marker GFP:SYP132 (Enami et al., 2009), and their fluorescence signal followed the membrane as it retracted inward upon plasmolysis (Figure 3c,d). These observations demonstrate that both MO25A proteins localize to the plasma membrane. The nuclear localization of MO25A1:RFP was further validated by co-staining with DAPI (Figure 3e). Together, these data indicate that, despite their genetic redundancy, MO25A1 and MO25A2 exhibit subtly distinct localization patterns, suggesting a degree of functional diversification.

**Figure 3.**
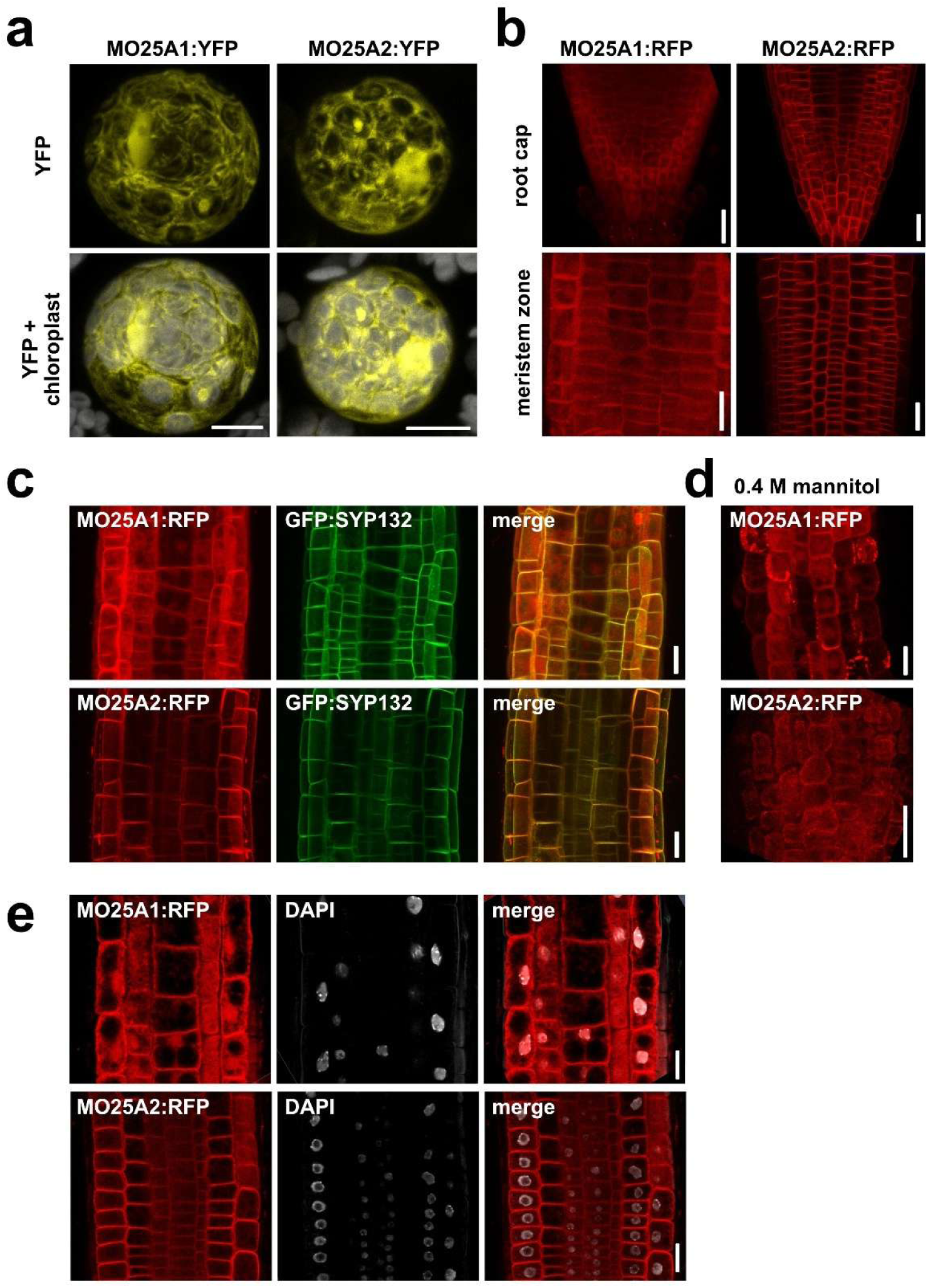
Subcellular localization of MO25A1 and MO25A2. a) Transient expression of MO25A1:YFP and MO25A2:YFP *Arabidopsis* mesophyll protoplasts. Scale bar = 5 µm. b) Localization of MO25A1:RFP and MO25A2:RFP in root cap and meristematic zone of stably transformed plants. c) Colocalization of MO25A1:RFP and MO25A2:RFP with GFP:SYP132. d) Localization of MO25A1:RFP and MO25A2:RFP in roots subjected to plasmolysis induced by treatment with 0.4 M mannitol for 60 min. and their colocalization with GFP:SYP132. e) Colocalization of MO25A1:RFP and MO25A2:RFP with DAPI-stained nuclei. Scale bars in b-e = 20 µm.

### MO25A1 recovers smg7-6 fertility by restoring TDM1 localization to M-bodies

To investigate the role of MO25A1 and MO25A2 in meiotic exit, we generated *mo25a1-1 smg7-6* and *mo25a2-1 smg7-6* double mutants and assessed their fertility. The *mo25a-1* mutation partially restored the fertility and pollen production in *smg7-6* plants, although the effect was less pronounced than that of the *mo25a-2* allele (Figure 4a,b). In contrast, the *mo25a2-1* mutation had an opposite effect, further reducing fertility and pollen production in the *smg7-6* background (Figure 4a,b). These results support the notion that the two proteins have undergone functional diversification and that suppression of the *smg7-6* phenotype is specific to MO25A1.

**Figure 4.**
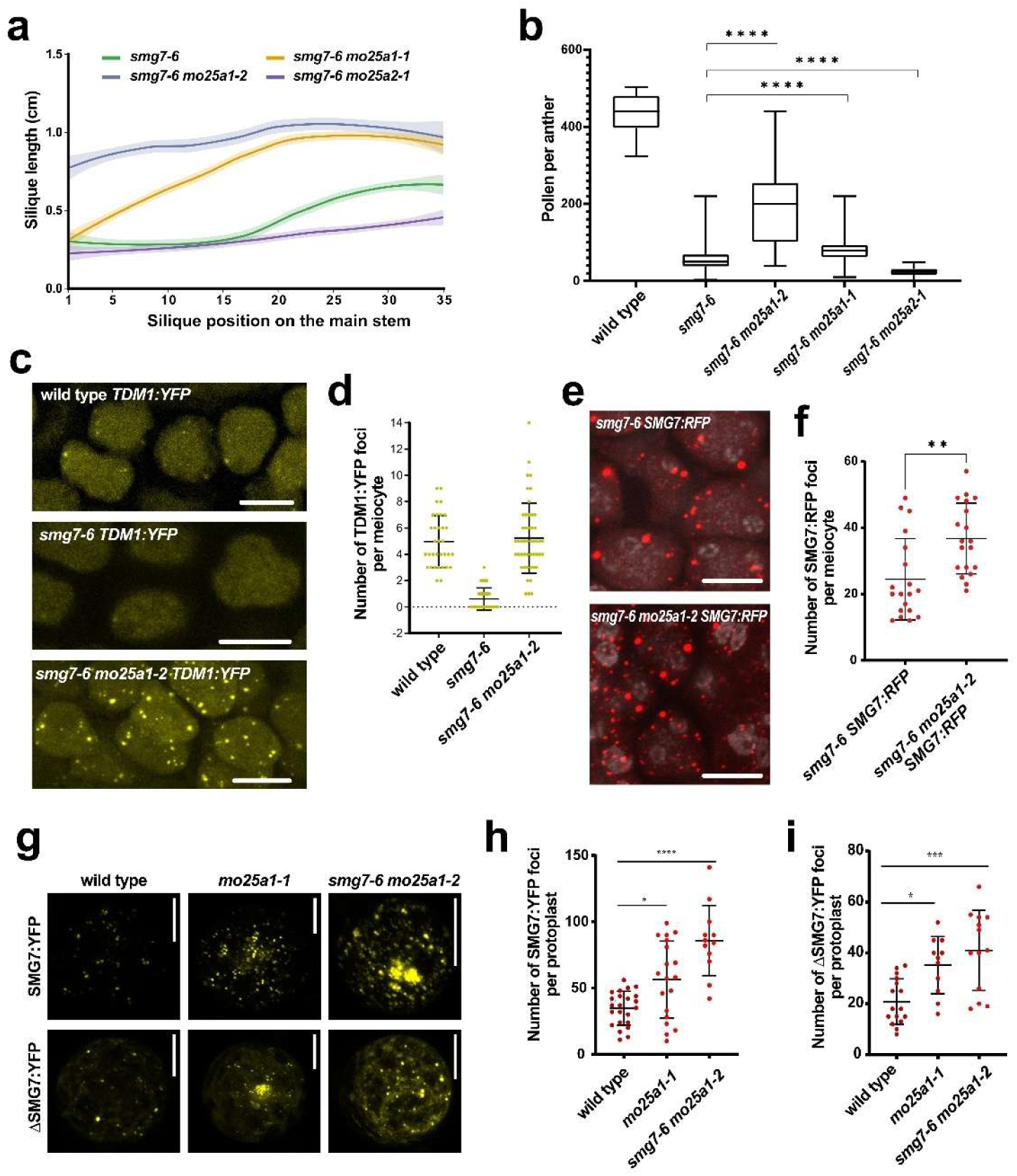
MO25A1 affects the partitioning of SMG7 and TDM1 into M-bodies. a) Quantification of silique length along the main inflorescence bolt in plants of the indicated genotypes; *smg7-6* (n = 18), *smg7-6 mo25a1-1* (n = 20), *smg7 mo25a1-2* (n = 22), and *smg7 mo25a2-1* (n = 3). Data trends were fitted using locally weighted scatterplot smoothing (LOESS, colored lines) and shaded area indicate 95% confidence intervals. b) Box-and-whisker plot showing the number of viable pollen per anther (**** P ˂ 0.0001, Kruskal Wallis with Dunn posthoc test, wild type n = 15, *smg7-6* n = 154, *smg7-6 mo25a1-2* n = 15, *smg7 mo25a1-1* n = 154, and *smg7 mo25a2-1* n = 67. c) Localization of TDM1:YFP in telophase II meiocytes of wild type and indicated mutants. d) Dot plots showing the average number of TDM1:YFP foci per telophase II meiocyte. e) Localization of SMG7:RFP in telophase II meiocytes of *smg7-6* and *smg7-6 mo25a1-2* plants. f) Dot plots showing the average number of SMG7:RFP foci per telophase II meiocyte (** P ˂ 0.01, Wilcoxon rank-sum test, *smg7-6 SMG:TagRFP* n = 19, *smg7-6 mo25a1-2* SMG7:TagRFP n = 19). g) Localization of SMG7:YFP and ΔSMG:YFP in transfected mesophyll protoplasts isolated from the indicated genotypes. h,i) Dot plots showing the average number of SMG7:YFP (h) and ΔSMG:YFP (i) foci in transfected mesophyll protoplasts (**** P ˂ 0.0001, *** P ˂ 0.001, * P ˂ 0.05, Kruskal Wallis test with post hoc Holm correction). Scale bars = 10 µm.

Reduced fertility in *smg7-6* plants is caused by insufficient recruitment of TDM1 to M-bodies by the truncated SMG7 protein (Cairo et al., 2022). Therefore, we next investigated whether *mo25a1-2* restores TDM1 localization to M-bodies in *smg7-6* mutants. Indeed, whereas number of TDM1 foci and their intensity were drastically reduced during telophase II in *smg7-6* meiocytes, the number of TDM1 foci was restored to wild type levels in *mo25a1-2 smg7-6* double mutants (Figure 4c,d). We also observed increased number of SMG7:RFP foci (Figure 4e,f) suggesting that MO25A1 may antagonize partitioning of SMG7 into M-bodies.

This observation raised the possibility that impaired MO25A1 function compensates for insufficient partitioning of the truncated SMG7 protein into condensates in *smg7-6* plants. To test this hypothesis, we analyzed the localization of SMG7:YFP and ΔSMG7:YFP, a truncated protein that mimics the *smg7-6* mutation, in mesophyll protoplasts. We observed significantly increased number of both SMG7:YFP and ΔSMG7:YFP foci in *smg7-6 mo25a1-2* protoplasts, while an intermediate effect was detected in the *mo25a1-1* mutant (Figure 4g-i). These data suggest that MO25A1 antagonizes the localization of SMG7 to P-bodies and M-bodies, and that loss of MO25A1 function enhances the fertility of *smg7-6* plants by improving the recruitment of ΔSMG7-TDM1 complexes to M-bodies.

### MO25A1 localizes to SGs and M-bodies under heat stress

The effect of MO25A1 on M-body formation raised the question of whether this protein physically associates with these structures. Under standard growth conditions, signals from both MO25A1:RFP and MO25A2:RFP expressed from their native promoters were undetectable in meiocytes, although fluorescence was readily observed in the surrounding tapetal cells (Figure 5a). MO25A1 was previously identified in a proteomic study as a component of heat-induced SGs (Kosmacz et al., 2019). Given that M-bodies possess an SG-like outer shell, we hypothesized that heat stress might promote the accumulation of MO25A proteins in these structures if they are expressed during meiosis. Indeed, exposure to 39°C induced the formation of distinct MO25A1 foci in telophase II meiocytes, while no comparable structures were observed for MO25A2 (Figure 5a).

**Figure 5.**
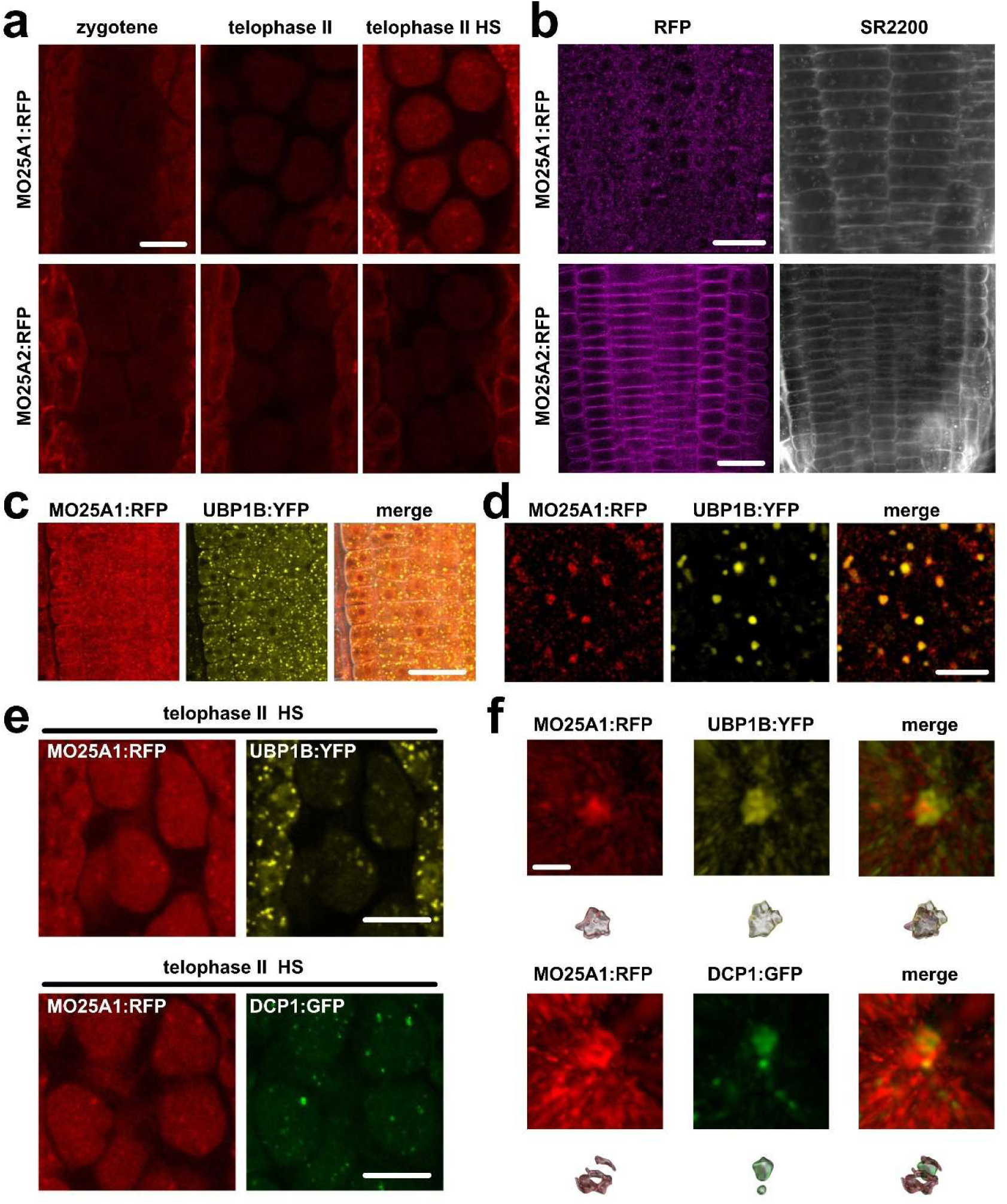
Localization of MO25A1 to stress granules and M-bodies. a) Localization of MO25A1:RFP and MO25A2:RFP in meiocytes under control conditions and folowing heat stress (HS, 39°C for 30 min). Scale bar = 10 µm. b) Localization of MO25A1:RFP and MO25A2:RFP in root meristem cells following heat stress. Scale bar = 20 µm. c,d) Colocalization of MO25A1:RFP with UBP1:YFP in root cells subjected to heat stress. Images are shown at low (c) and high (d) magnification. Scale bars = 20 µm (c) and 5 µm (d). e) Colocalization of MO25A1:RFP with the SG marker UBP1B:YFP and P-body DCP1:GFP marker in heat-stressed telophase II meiocytes. Scale bar = 10 µm. f) Super-resolution micrographs of the indicated protein condensates displayed by 3D view (top panels) and 3D rendering (bottom panels) generated using Imaris software. Scale bar = 1 μm.

To determine whether this difference reflects expression in meiocytes or a difference in the ability of the two proteins to partition into SGs, we also examined their localization in roots. Whereas MO25A2 remained predominantly associated with the plasma membrane, MO25A1 formed prominent cytoplasmic granules upon heat treatment (Figure 5b). These granules co-localized with the stress granule marker UBP1B:YFP (Figure 5c,d), confirming their identity as SGs. Together, these results suggest that MO25A1, but not MO25A2, has an inherent propensity to partition into SG-like condensates.

The MO25A1 signal in meiocytes overlapped with both UBP1B:YFP and DCP1:GFP, the latter serving as a marker of P-bodies (Figure 5e). A more detailed analysis using structured illumination microscopy revealed that MO25A1 was primarily positioned adjacent to DCP1, while showing more extensive overlap with UBP1B (Figure 5f). These findings indicate that MO25A1 associates with the SG-like shell of M-bodies and that this association becomes detectable under heat stress. Given that the SG-like shell is also present in M-bodies under standard growth conditions, it is possible that MO25A1 is constitutively associated with these structures, albeit at levels below the detection limit.

### MO25A1 and MO25A2 interact with condensate-associated CIPK kinases

MO25 is an evolutionary conserved scaffold protein that regulates STE20-like kinases in both yeast and animals. Therefore, we hypothesized that, similarly in plants, MO25 functions through the regulation of an associate kinase. In Arabidopsis, STE20-like kinases are represented by the MAP4K kinase family (Champion et al., 2004). The closest homolog of the human STE20-like kinase MST3 is Arabidopsis MAP4K1 (At1g53165). Structural studies of the human MST3-MO25β complex identified the interaction interphase between these proteins (Mehellou et al., 2013), and this region is highly conserved in MAP4K1 (Figure S2). Yeast two-hybrid assay confirmed that MAP4K1 and its close homolog MAP4K2 interact with MO25B, whereas no interaction was detected with MO25A proteins (Figure 6a, Figure S3). These data indicate that although the MAP4K1-MO25B represents an evolutionary conserved regulatory module, MO25As have diverged in plants and likely interact with other kinases.

**Figure 6.**
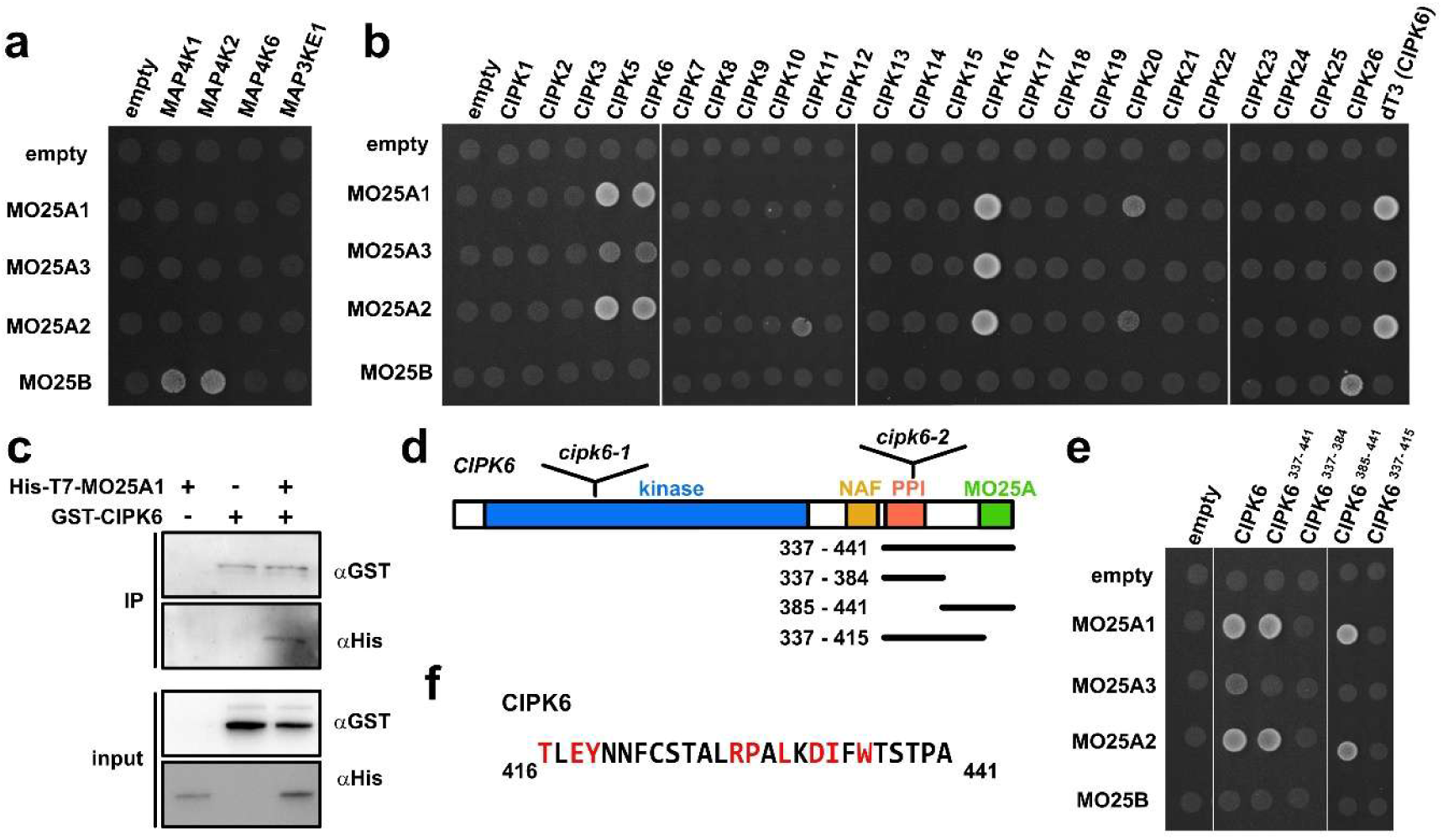
Identification of MO25-interacting kinases. a,b) Interaction of MO25 proteins cloned in pGBKT7 bait vector with MAPK4K (a) and CIPK (b) kinases cloned into the pGAD4242 and pGADT7 DEST pray vectors, respectively, assessed by yeast two-hybrid assay. Yeast colonies were grown on SD-HLT plates supplemented with 1 mM 3AT. dT3 represents a CIPK6 fragment identified in the yeast two-hybrid screen. c) Western blot analysis of an *in vitro* pull-down assay using His-T7-MO25A1 and GST-CIPK6 proteins bound to glutathione Sepharose 4B. Proteins were detected using ant-His and anti-GST antibodies. d) Schematic representation of the domain architecture of CIPK6. The positions of T-DNA insertions in *cipk6* alleles and regions used to map the MO25A1-intereacting domain are indicated. e) Interaction of MO25 proteins with truncated CIPK6 variants assessed by yeast two-hybrid assay. Yeast colonies were grown on SD-HLT plates supplemented with 1 mM 3-AT. f) Amino acid sequence of the C-terminal region of CIPK6. Residues conserved among MO25- interacting CIPKs are highlighted in red.

To identify these kinases, we performed an yeast two-hybrid screen of Arabidopsis cDNA libraries using MO25A1 as a bait. Among the five candidate interactors identified, the most prevalent hits were the CBL-interacting protein kinases CIPK6, CIPK20, and CIPK5 (Table S1). The Arabidopsis CIPK family comprises 26 plant-specific kinases that associate with CBL calcium sensors to decode calcium signals and regulate ion homeostasis, stress responses, and plant development (Tang et al., 2020; Dong et al., 2021). We therefore systematically tested all Arabidopsis CIPKs, with the exception of CIPK4, for interaction with MO25 proteins. Both MO25A1 and MO25A2 associated with CIPK5, CIPK6, CIPK16, and CIPK20, whereas MO25A3 interacted with CIPK5, CIPK6, and CIPK16 (Figure 6b and Figure S3). In addition, MO25A2 showed weak binding with CIPK11. In contrast, MO25B associated exclusively with CIPK26. We further validated the MO25A1-CIPK6 interaction by in vitro immunoprecipitation (Figure 6c), and confirmed most of MO25A1 and MO25A2 associations by bimolecular fluorescence complementation (BiFC; Figure S4a). MO25A3 produced no or only weak BiFC signals (Figure S4a), consistent with its weaker associations detected in the yeast two-hybrid assay (Figure 6b). These results demonstrate that MO25A and MO25B proteins exhibit distinct kinase binding profiles and suggest that MO25A isoforms have evolved as regulators of a subset of CIPKs.

Given the specific association of MO25A proteins with a subset of CIPKs, we next sought to identify the CIPK region responsible for this binding. CIPKs consist of an N-terminal kinase domain and a C-terminal regulatory region containing the CBL-interacting NAF motif and a PP2C-interacting motif (PPI) implicated in phosphatase-mediated regulation (Figure 6d). We focused on CIPK6 because it was the strongest candidate recovered in the yeast two-hybrid screen, being represented by several independent cDNA clones (Table S1). Notably, the shortest cDNA clone encoded the C-terminal region beginning at the PPI motif. Yeast two-hybrid assays using fragments spanning this region narrowed the site required for MO25A binding to the last 26 amino acids of CIPK6 (Figure 6d,e).

Sequence analysis of the C-termini identified a conserved motif that was present in all MO25A1- associated kinases, with the exception of CIPK11 (Figure 6f and Figure S5). Inspection of the remaining Arabidopsis CIPKs revealed the same motif in CIPK2, CIPK10, and CIPK25. Consistently, both CIPK10 and CIPK25 were found to associate with MO25A proteins in BiFC assays. Together, these findings indicate that the specificity of MO25A proteins toward selected CIPK kinases is largely determined by a conserved motif located at the extreme C-terminus.

To gain further insight into the MO25A1-associated CIPKs, we analyzed their subcellular localization in transiently transfected protoplasts. CIPK5, CIPK6, and CIPK20 localized to cytoplasmic granules, whereas CIPK16 was restricted to the nucleus and nuclear granules (Figure 7a). CIPK6 and CIPK20 colocalized with DCP1, suggesting their association with P-bodies, whereas CIPK5 showed no overlap with DCP1 (Figure 7b). We therefore examined whether CIPK5 associates with stress granules (SGs). Upon heat stress, the CIPK5 signal reorganized into smaller, more abundant foci that colocalized with the SG marker G3BP-2 (Figure 7c). Finally, CIPK16 nuclear foci colocalized with COILIN, suggesting an association with Cajal bodies. Thus, the CIPKs that most consistently interacted with MO25A1 shared a propensity to localize to distinct RNP condensates.

**Figure 7.**
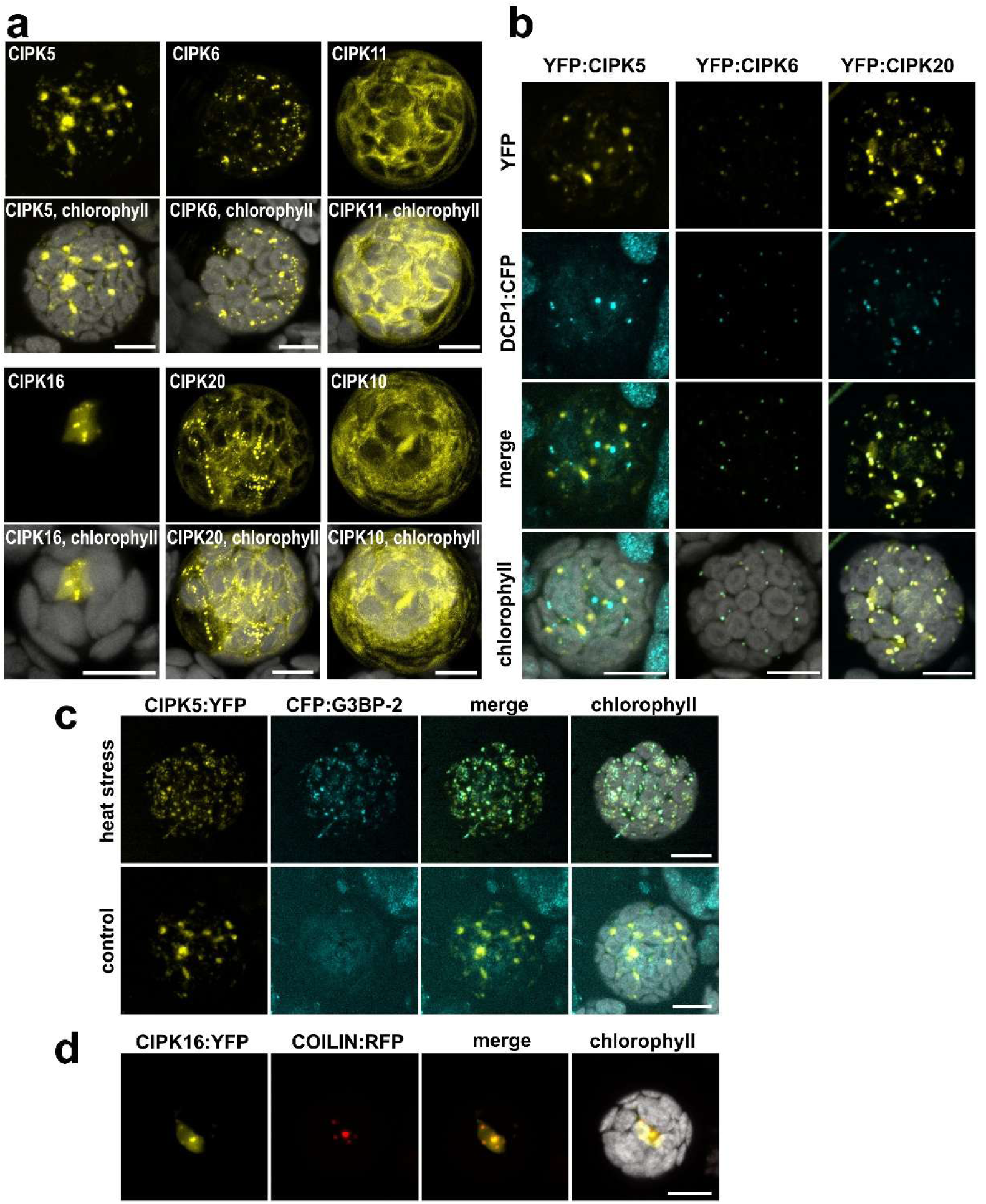
Localization of MO25A-interacting CIPKs in protoplasts. a) Localization of the indicated YFP-fused CIPKs in protoplasts. b) Colocalization of CIPK5, CIKP6, and CIPK20 with the P-body marker DCP1. c) Colocalization of CIPK5 with the SG marker G3BP-2 under control and heat stress conditions. d) Colocalization of CIPK16 with the Cajal body marker COILIN. Scale bars = 10 µm.

### CIPK6 is required for pollen development and affects partitioning of SMG7 to P-bodies

Given their localization to P-bodies, which constitute the core of M-bodies and recruit TDM1 (Cairo et al., 2026), we next focused on CIPK6 and CIPK20. Plants carrying a T-DNA insertion in *CIPK20* displayed no obvious defects in growth or fertility under standard conditions (Figure 8b). In contrast, disruption of *CIPK6* by a T-DNA insertion within the conserved kinase domain (*cipk6-1*) resulted in severe growth retardation and sterility (Figure 8a). To our knowledge, such a severe phenotype has not been reported for previously characterized *cipk6* mutants harboring more C-terminal T-DNA insertions (Tripathi et al., 2009; Held et al., 2011). To verify that defects were caused by loss of CIPK6 function, we complemented *cipk6-1* with a genomic construct spanning the *CIPK6* locus. Examination of anthers revealed a drastic reduction in pollen production in *cipk6-1* plants (Figure 8b,c). Notably, a significant decrease in pollen number was also observed in otherwise phenotypically normal *cipk6- 1/+* heterozygotes. Analysis of postmeiotic cells in *cipk6-1* anthers revealed, in addition to normal tetrads, various abnormal meiotic products (Figure 8d,e). These included polyads, which are typically indicative of aberrant chromosome segregation during meiosis, as observed for example in *smg7-6* mutants. We also detected atypical structures containing elongated cells within tetrads, or their linear arrangement instead of characteristic tetrahedral organization, indicating abnormalities in the patterning of meiotic division (Figure 8d,e).

**Figure 8.**
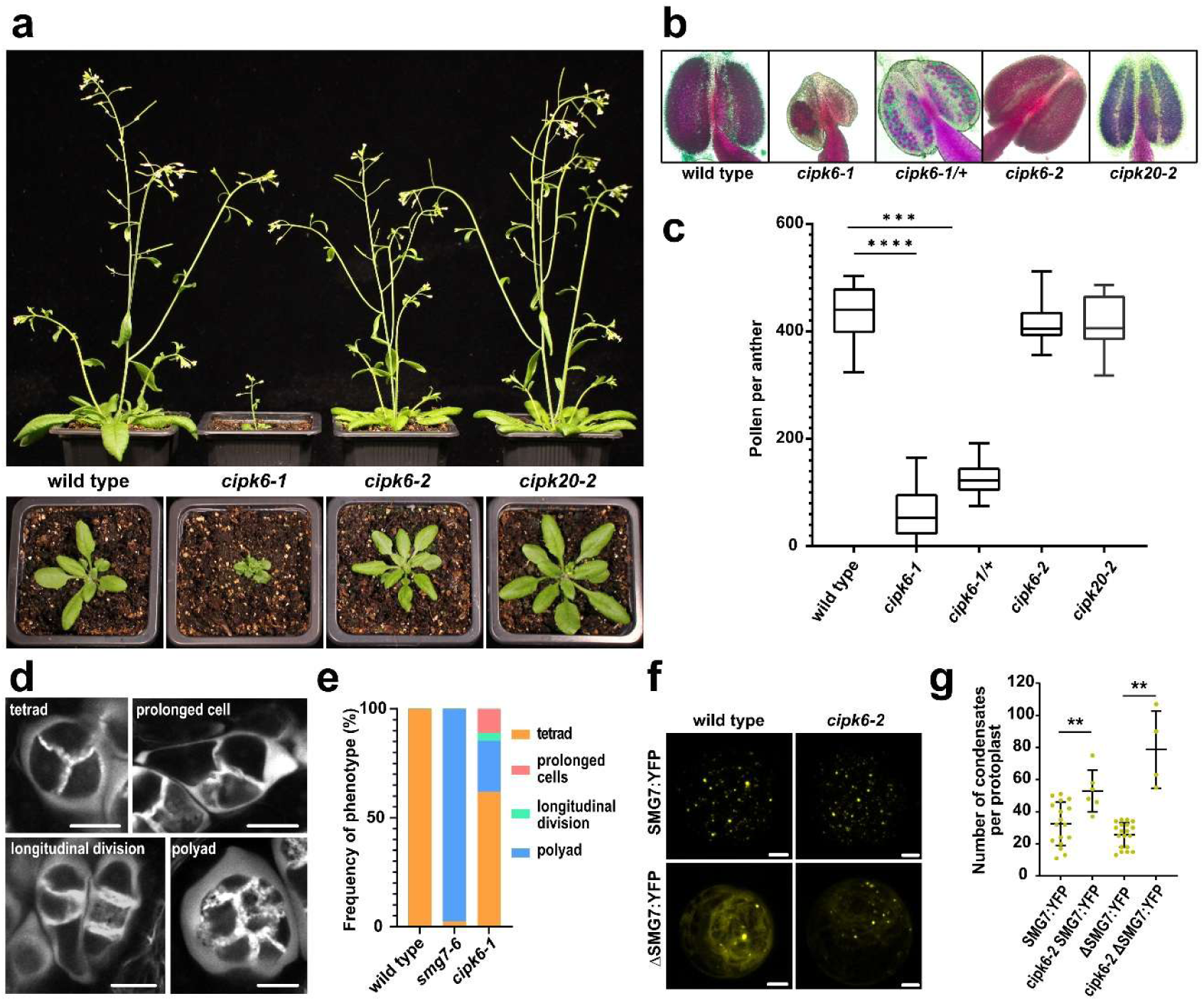
Characterization of Arabidopsis *cipk6* mutants. a) Representative phenotypes of 6-week-old plants (top panel) and 3-week-old rosettes (bottom panel) of the indicated genotypes. b) Anthers of the indicated genotypes after Alexander staining. c) Box-and-whisker plot showing the number of viable pollen per anther (**** P ˂ 0.0001, *** P ˂ 0.001 Kruskal Wallis with Dunn posthoc test, wild type n = 15*, cipk6-1* n = 35, *cipk6-1/+* n = 28, *cipk6-2* n = 10, and *cipk20-2* n = 7). d) Tetrads and aberrant meiotic products in *cipk6-1* mutants visualized by callose staining with SR2200. e) Quantification of meiotic tetrads and aberrant meiotic products in wild type (n = 676), *smg7-6* (n = 254), and *cipk6-1* (n = 211) plants. f) Localization of SMG7:YFP and ΔSMG:YFP in transfected mesophyll protoplasts isolated from the indicated genotypes. g) Dot plots showing the average number of SMG7:YFP foci in transfected mesophyll protoplasts from wild type and *cipk6-2* plants (** P ˂ 0.01, Test - Mann–Whitney U test). Scale bars = 10 µm.

These observations strongly indicate that CIPK6 is required for the proper completion of meiosis. Together with its localization to P-bodies, this finding raised the possibility that MO25A1 exerts its meiotic function through the regulation of CIPK6. To investigate this hypothesis, we took advantage of a previously described allele carrying a T-DNA insertion within the C-terminal PPI domain (here designated *cipk6-2*)(Held et al., 2011), which lacks the MO25-binding region. Although *cipk6-2* mutants are phenotypically similar to wild type under standard growth conditions and remain fully fertile (Figure 8a), we asked whether the mutation affects SMG7 localization in a manner similar to *mo25a1-2*. Indeed, localization studies in transiently transfected protoplasts showed that *cipk6-2* phenocopies the *mo25a1-2* mutation by enhancing the partitioning of both SMG7:YFP and ΔSMG7 into condensates (Figure 8f,g). Together, these results identify CIPK6 as a possible downstream effector of MO25A1 in meiosis.

## Discussion

MO25 proteins are highly conserved across eukaryotes and have been extensively characterized in animals and yeast, where they function as allosteric activators of several kinases in the STE20 superfamily (Boudeau et al., 2003; Nelson et al., 2003; Kanai et al., 2005; Filippi et al., 2011). MO25 homologues are also present in plants, where they diverged at the base of land plants into two distinct clades, MO25A and MO25B (Bizotto et al., 2018). Here, we identified kinases that interact with Arabidopsis MO25 proteins and are presumably regulated by them. Similar to their counterparts in yeast and mammals, plant MO25B interacts with MAP4K kinases, which are homologous to the MST kinases of the STE20 family, indicating that the MO25–MST/MAP4K regulatory module is evolutionarily conserved. In contrast, the MO25A clade appears to have acquired specificity for plant-specific CIPKs, revealing a lineage-specific diversification of MO25–kinase interactions.

CIPKs are Ser/Thr protein kinases that interact with calcium sensors of the CBL (calcineurin B-like) family. CBL–CIPK complexes decode changes in intracellular Ca²⁺ levels and regulate diverse targets involved in nutrient homeostasis, ion transport, and responses to abiotic and biotic stresses (Tang et al., 2020; Dong et al., 2021). CIPKs interact with CBLs through the NAF motif in their C-terminal regulatory region, which promotes kinase activation and/or recruitment to specific subcellular compartments and substrates (Albrecht et al., 2001; Guo et al., 2001). A second regulatory module in the C-terminus, the PPI motif, mediates interaction with PP2C-type phosphatases, which can antagonize CIPK signaling by dephosphorylating CIPKs or their downstream substrates (Ohta et al., 2003; Lee et al., 2007; Lan et al., 2011). Together, these modular interactions and the combinatorial pairing of different CIPKs with CBLs and PP2C phosphatases generate a versatile signaling network that enables plants to tailor responses to diverse environmental and cellular cues.

MO25A may represent an additional regulatory component of this network. We found that MO25A interacts with a subset of CIPKs through their extreme C-terminus, with a specificity that appears to be defined by a conserved sequence motif. Whether and how MO25A affects CIPK activity remains to be determined. By analogy to the allosteric activation of animal MST3 by MO25 (Mehellou et al., 2013), MO25A could directly activate CIPKs by inducing conformational changes in the kinase domain. Alternatively, MO25A may modulate CIPK activity indirectly by influencing the interaction of CIPKs with CBLs or PP2C phosphatases. MO25A could also regulate CIPK function by controlling their subcellular localization, as MO25 has been shown to mediate recruitment of the Sid2 kinase to the cell division site during cytokinesis in fission yeast (Ye et al., 2026).

Studies in rice and the moss *Physcomitrium patens* (Ta et al., 2023), together with our analysis in *Arabidopsis*, establish that MO25A function is essential in plants. MO25A proteins have undergone independent duplication events in diverse plant lineages (Bizotto et al., 2018). In rice, the *OsMO25A1* paralogue is essential, whereas *OsMO25A2* expression appears to be restricted to mature pollen. In contrast, the *P. patens* homologues *PpMO25A1* and *PpMO25A2* are nearly identical, and inactivation of either gene alone is viable, whereas simultaneous inactivation is lethal (Ta et al., 2023).

*Arabidopsis* contains three MO25A paralogues. MO25A1 and MO25A2 arose through a duplication in the Brassicaceae and their function is essential for viability, whereas MO25A3 is evolutionarily more distant and originated through a retrotransposition event (Bizotto et al., 2018). This evolutionary divergence is also reflected in the weaker interaction of MO25A3 with CIPKs, suggesting that its binding specificity may have diverged. Although MO25A1 and MO25A2 interact similarly with CIPKs and are largely redundant, our study reveals evidence of partial functional diversification. Their inactivation has opposite effects on the fertility of *smg7-6* mutants, and they also exhibit distinct subcellular localization patterns. In addition to its association with the plasma membrane, MO25A1 shows broader subcellular localization, including in the nucleus, and has a propensity to relocalize to SGs upon heat stress, whereas MO25A2 remains restricted to the plasma membrane. This partial diversification may therefore add another layer of complexity to CIPK regulation, potentially allowing individual CIPKs to be regulated differently depending on the MO25A paralogue and cellular context.

Molecular functions of CIPKs are traditionally associated with membrane-localized processes that regulate transporter activity and ion homeostasis (Tang et al., 2020; Dong et al., 2021). However, several CIPKs also have established non-membrane functions. These include CIPK6- and CIPK11- mediated regulation of transcription factors involved in abscisic acid signaling and iron-starvation responses (Zhou et al., 2015; Gratz et al., 2019; Mao et al., 2026), the role of CIPK20 in controlling microtubule dynamics during drought-induced stomatal closure (Li et al., 2024), and CIPK6- dependent oxidative stress responses mediated through the cytosolic protein Rd2 (Gutierrez-Beltran et al., 2017). Notably, the CIPKs displaying the strongest interactions with MO25A1/2 were associated with RNP condensates in the protoplast transient expression system. Specifically, CIPK6 and CIPK20 localized to P-bodies, CIPK5 to stress granules, and CIPK16 to Cajal bodies. These observations suggest that the MO25A-CIPK module may participate in the regulation of RNP condensate dynamics and post-transcriptional gene expression.

Indeed, we identified MO25A1 in a suppressor screen for increased fertility of *smg7-6* mutants. The *mo25a1-2* allele appears to enhance partitioning of SMG7 into condensates, thereby compensating for the inefficient recruitment of TDM1 into M-bodies by the truncated SMG7 protein in *smg7-6* mutants. Although it remains unclear whether this function is mediated through a CIPK, CIPK6 emerges as a plausible candidate. *cipk6-1* mutants, carrying a disruption in the N-terminal kinase domain, exhibit stunted growth, reduced fertility, and a high frequency of abnormal meiotic products indicative of defects in meiotic division. Moreover, the *cipk6-2* allele, which produces a truncated CIPK6 lacking the C-terminal MO25-binding domain, enhances SMG7 condensation similarly to MO25A1 deficiency. Together, these observations suggest that MO25A1 and CIPK6 may function in a common pathway controlling SMG7 condensation and M-body organization.

In conclusion, our work uncovers interaction between the MO25 scaffold protein and CIPK kinases and suggests that MO25A–CIPK complexes form a regulatory module with potential roles in the organization and function of diverse RNP condensates.

## Material and methods

### Plant material and growth conditions

*Arabidopsis thaliana* (ecotype Col-0), mutant lines and transgenic lines were grown on soil in growth chambers at 21°C at 50-60% of humidity under 16/8 h light/dark cycles. Roots were analysed from 4 days old seedlings grown in agar plates (0.8% w/v, pH 5.7) supplemented with half-strength Murashige and Skoog medium (½ MS). The following *Arabidopsis* mutant and reporter lines were used in this study: *smg7-6* (Riehs-Kearnan et al., 2012), *GFP-SYP132* (Enami et al., 2009), *TDM1:YFP*(Cairo et al., 2022) and *SMG7:TagRFP* (Cairo et al., 2022), *UBP1b:YFP* and *DCP1:G3GFP* (Cairo et al., 2026). The mutant lines *mo25a1-1* (SALK_073939)*, mo25a2-1* (SALK_076387)*, cipk6-1* (GABI_201C06)*, cipk6-2* (GABI_448C12), and *cipk20-2* (GABI_533G02) were obtained from the Nottingham Arabidopsis Stock Centre. All the mutants used in the study were PCR genotyped to confirm the presence of T-DNA insertions using primer indicated in Table S2. *MO25A1:TagRFP* and *MO25A2:TagRFP* reporter lines were generated in this study.

### Suppressor screen

The suppressor screen for mutations restoring fertility in *smg7-6* mutant plants was performed as previously described (Capitao et al., 2021). De novo mutations associated with the phenotype were identified by whole-genome sequencing and mapped using ArtMAP (Javorka et al., 2019). The EMS225 mutant was genotyped using the derived cleaved amplified polymorphic sequence (dCAPS) method with the primers chr4_9678061_FW and chr4_9678061_Rev (Table S2).

### RNA extraction and RT PCR

Total RNA was isolated from seedlings using RNA Blue (TOP-Bio). Residual genomic DNA contamination was eliminated by treating the isolated RNA with DNase I (Roche). For first-strand cDNA synthesis, 2.5 μg of the treated total RNA was reverse transcribed using M-MLV Reverse Transcriptase (Invitrogen) and primed with oligo(dT) primers. To analyze the transcripts in the *EMS225* mutant, PCR amplification was performed using the gene-specific primers AT4G17270_2FW and AT4G17270_2Rev. The resulting PCR products were sequenced.

### Fertility assays

Pollen count and viability were determined using Alexander staining, described previously (Alexander, 1969), and imaged with an Axioscope.A1 transmitted light microscope (Zeiss, 20×/0.5 objective), equipped with an Axiocam 105 camera and Visiview software (Visitron Systems). Meiosis was assessed by DAPI for DNA staining and Renaissance 2200 (SR2200) for callose staining in whole anthers as described (Capitao et al., 2021), and imaged on a LSM780 confocal microscope (Zeiss, C Plan-Apochromat 63x/1.4 Oil DIC M27). Silique length was assessed from images of main stems scanned by an Epson scanner and the siliques were measured using the Fiji Analyze/Measure function (Schindelin et al., 2012).

### Plasmid construction

To generate *MO25A1::MO25A1-TagRFP* and *MO25A2::MO25A2-TagRFP* constructs, genomic fragments containing the putative promoters and the ORFs of genes were amplified with the primers AT4G17270_FW_GC and AT4G17270_2Rev_GC, AT5G47540_FW_GC and AT5G47540_Rev_GC, respectively. The fragments were introduced into pENTR™/D-TOPO (Invitrogen) using blunt-end TOPO® Cloning reactions, and after LR recombination into pGWB659 vector. The final constructs were introduced into mutant lines via *Agrobacterium tumefaciens-*mediated transformation using the floral dip method. For genetic complementation of *EMS225* line, the *MO25A1* gene was amplified with the primers AT4G17270_FW_GC and AT4G17270_Rev_GC. The PCR product was cloned into pENTR™/D-TOPO™ and recombined into pGWB601.

For the yeast two hybrid constructs, MO25A1, MO25A2, MO25A3, MO25B, cDNAs were amplified using the primers AT4G17270_FW, AT4G17270_Rev, AT5G47540_FW, AT5G47540_Rev, AT5G18940_FW, AT5G18940_Rev, AT2G03410_FW and AT2G03410_Rev (Table 2). PCR fragments were cleaved with restriction enzymes *Sma*I and *Pst*I and cloned into the pGBKT7 (bait) vector. CIPK kinases were amplified using the corresponding pairs of primers (Table S2) and transferred into pENTR™/D-TOPO. The coding sequences of CIPK kinases were subcloned from their entry clones into the destination vectors pGADT7-DEST (pray) using LR Clonase™ (Invitrogen). For MAP4K1, MAP4K2, MAP4K6 PCR fragments were digested with EcoRI and SalI and the resulting fragments were cloned into pGAD424 (pray) vector. cDNA of MAP3KE1 was amplified using AT3G13530_FW and AT3G13530_Rev primers and digested with *BamHI* and *BglII* restriction enzymes before cloning into the pGAD424 (pray) vector. To generate deleted version of CIPK6 protein, we used the method described in (Stoynova et al., 2004).

For yeast two-hybrid assays, the cDNAs of MO25A1, MO25A2, MO25A3, and MO25B were amplified using the primers AT4G17270_FW/AT4G17270_Rev, AT5G47540_FW/AT5G47540_Rev, AT5G18940_FW/AT5G18940_Rev, and AT2G03410_FW/AT2G03410_Rev, respectively (Table S2). The PCR products were digested with SmaI and PstI and cloned into the bait vector pGBKT7. CIPK kinase coding sequences were amplified using the corresponding primer pairs (Table S2) and cloned into pENTR™/D-TOPO™. The CIPK coding sequences were subsequently transferred from the entry clones into the prey vector pGADT7-DEST by LR recombination using LR Clonase™ (Invitrogen). For MAP4K1, MAP4K2, and MAP4K6, PCR products were digested with EcoRI and SalI and cloned into the prey vector pGAD424. The cDNA of MAP3KE1 was amplified using the primers AT3G13530_FW and AT3G13530_Rev and digested with BamHI and BglII prior to cloning into the prey vector pGAD424. To generate a deletion variant of the CIPK6 protein, the method described by Stonyova et al. (Stoynova et al., 2004) was used.

The expression constructs for transient assays in protoplasts were generated using the Gateway® cloning system. Coding sequences of the target genes were amplified and initially cloned into the pENTR™/D-TOPO entry vector. These entry clones were then transferred into suitable Gateway destination vectors through LR recombination to produce fluorescent protein fusions. For subcellular localization analyses, constructs were introduced into pGWB441 and pGWB442, enabling the generation of C-terminal and N-terminal YFP-tagged proteins, respectively. For colocalization experiments, constructs were recombined into pGWB460 to obtain RFP-tagged fusion proteins.

Bimolecular Fluorescence Complementation (BiFC) assays were performed by recombining entry clones into the split-YFP destination vectors pGWcY, pcYGW, pnYGW, and pGWnY, allowing both N- and C-terminal fusion configurations for protein–protein interaction studies. The following constructs were used as subcellular markers: TDM1:YFP, DCP1:CFP, SMG7:YFP, and the ΔSMG7 (*smg7-6*) (Cairo et al., 2022), CFP:G3BP-2 (Schindfessel et al., 2026), COILIN:RFP (Fulneckova et al., 2026).

Constructs for *MO25A1* and *CIPK6* expression in E. coli were prepared as follows: the cDNA of MO25A1 and CIPK6 were amplified from total cDNA with the primers MO25-1-BAMHI-F and MO25-1- XHO1nostop-R, and CIPK6-BAMHI-F and CIPK6-XHOI-STOP-R respectively. Subsequently, the cDNA of *MO25A1* was digested with the restriction enzymes BamHI and XhoI, and cloned into the respective restriction sites of the vector pET28a, obtaining the construct pET28-His-T7-Mo25. The cDNA of *CIPK6* was digested with the restriction enzymes BamHI and XhoI, and cloned into the respective restriction sites of the vector pGEX4T-1, obtaining the construct pGEX4T-1-GST-CIPK6.

### Protein pull-down and western

pET28-His-T7-Mo25 and pGEX4T-1-GST-CIPK6 were expressed in *E. coli* Rossetta (DE) cells (Novagen). The Histidine-tagged protein was purified using the His Mag Sepharose™ Ni Magnetic Beads (Cytiva) according to the manufacturer’s instructions. To purify the protein GST-CIPK6, the extract was incubated with Glutathione Sepharose 4B (GE Healthcare Life Sciences), and purified according to the manufacturer’s instructions. The purified proteins were subsequently dialyzed into storage buffer (20 mM Tris-HCl pH 7.4, 50 mM NaCl), and concentrated via ultrafiltration (Amicon® Ultra-0,5 [Merck Millipore]) prior storage at -80°C. The MO25A1-CIPK6 interaction assay was performed by incubating 0.5 µM of purified HIS-T7-MO25 and 0.5 µM of purified GST-CIPK6 in Binding buffer (PBS pH 7.3) for 2 hours at 25°C, following incubation with Glutathione Sepharose 4B (GE Healthcare Life Sciences) for 1.5 hours at 25°C. After 3 washes in Binding buffer, proteins were eluted from the beads by adding 1x Leammli sample buffer. For Western blotting, proteins were detected using mouse monoclonal antibody [HIS.H8] for 6x His tag (Abcam), or GST Tag rabbit polyclonal antibody (ThermoFisher A-5800).

### Yeast two-hybrid screening

The yeast two hybrid experiments were performed using the Matchmaker™ GAL4-based two-hybrid system (Clontech). The screen for identification of MO25A1 interacting partners was performed using the MO25A1 cDNA subcloned into pGBKT7. Bait vector was transformed into *Saccharomyces cerevisiae strain Y187* and mated with *AH190* strain harboring two different *Arabidopsis* cDNA libraries (oligo dT-primed and random-primed cDNA libraries, gift of the Hans Sommer Laboratory, Max Planck Institut, Germany). Resultant zygotes were selected by culturing on SD−Leu−Trp-His + 2,5 mM 3-amino-1,2,4-aminotriazole (3AT) plate for 7 days at 30°C. Plasmids collected from positive clones were then sequenced using the pGAD_FW and pGAD_Rev primers (Table S2).

For pairwise interaction assays, bait and pray vectors were transformed into Y187 and AH190 yeast strains respectively. Interactions between proteins were assayed by the mating method. Yeast cells carrying both plasmids were selected on SD-Leu -Trp medium and protein-protein interactions were screened on SD-Leu-Trp-His + 3AT medium and SD-Leu-Trp-His-Ade + 3AT medium. Empty vectors were used as negative controls for auto-activation. All experiments were performed in two independent biological replicates, each with three technical replicates.

### Microscopy

Arabidopsis mesophyll protoplast transfection was performed as previously described (Yoo et al., 2007). Imaging was carried out 16 h after transfection using a Zeiss LSM780 confocal microscope equipped with an LCI Plan-Neofluar 63×/1.3 Imm Korr DIC M27 objective. For heat-shock treatments, transfected protoplasts were incubated at 39 °C for 30 min in a heat block prior to imaging.

Protein localization in Arabidopsis roots was analyzed in 4-day-old seedlings. Seedlings were fixed in 4% formaldehyde in fixation buffer (FB; 1 mM EDTA, 0.1% Triton X-100 in 1× PBS, pH 7.0), vacuum-infiltrated for 15 min, incubated for an additional 45 min, and washed three times with FB. Cell walls were stained with SR2200 (1:1000 dilution in FB) for 30 min, followed by three washes with FB. DNA was stained with 5 µg mL⁻¹ DAPI in FB for 1 h. Samples were mounted in VECTASHIELD® Antifade Mounting Medium and imaged using a Zeiss LSM780 confocal microscope. Membrane association of fluorescently tagged proteins was assessed by mannitol-induced plasmolysis. Seedlings were incubated in 0.4 M mannitol for 1 h, and fluorescence redistribution was analyzed using a Zeiss LSM780 confocal microscope. Super-resolution imaging was performed using a ZEISS Elyra 7 microscope with lattice structured illumination microscopy (Lattice SIM), a Plan-Apochromat 63×/1.4 Oil DIC M27 objective, and two PCO Edge sCMOS cameras. Heat shock was applied by immersing sealed plates in a 39 °C water bath for 30 min, after which roots were rapidly fixed. Protein localization in *Arabidopsis* anthers was performed as described (Capitao et al., 2021).

### Image processing and analysis

ZEN software (blue and black edition) was used to process images obtained by laser scanning confocal microscopes. The 3D segmentation and the signal quantification of RNP granules from roots and meiocytes was performed with the Imaris 10.2.0 microscopy image analysis software, utilizing the volume rendering. The super-resolution images obtained with the microscope ZEISS Elyra 7 microscope were processed in ZEN Black 3.0 SR (Zeiss), utilizing the 3D SIM2 algorithm, with output sampling = 2 and scaled to raw.

## Acknowledgements

This work was supported by the Czech Science Foundation (22-31712S). Microscopy was performed at the Imaging Facility of the IEB AS CR and the core facility of CEITEC Masaryk University CELLIM funded by MEYS CR (LM2023050 Czech-BioImaging). We further acknowledge support of Plant Science Core Facility with plant cultivation.

