## Supplementary Material for "MO25 binds CBL-interacting protein kinases associated with ribonucleoprotein condensates and regulates meiotic exit"

Figures S1 – S5

Table S1

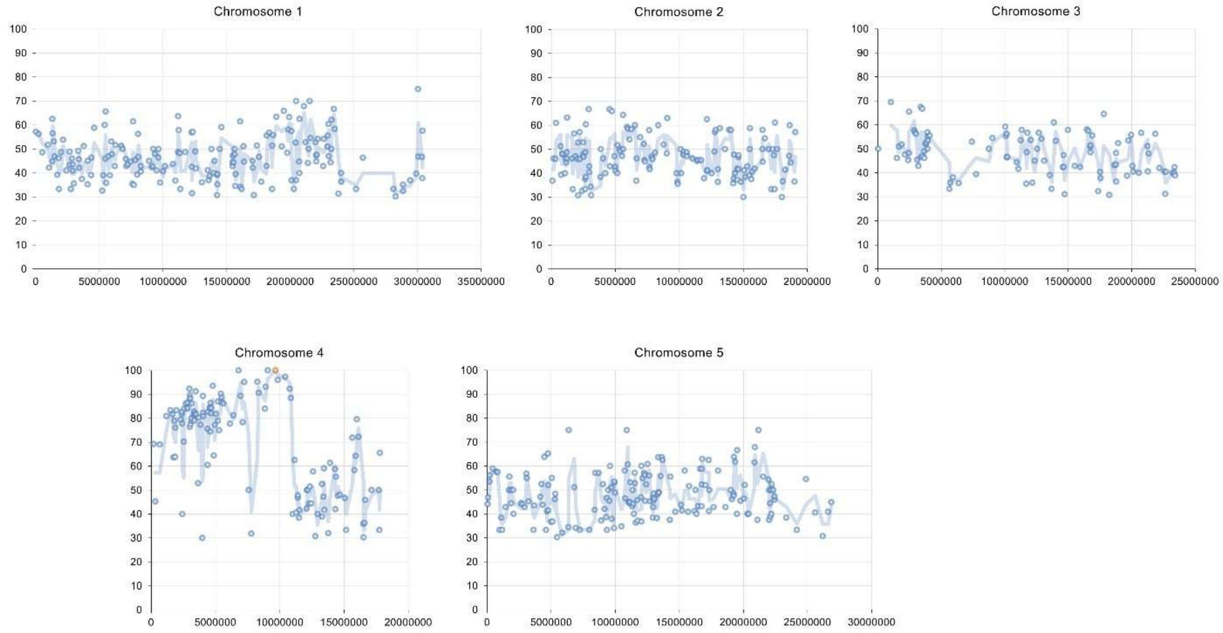


**Figure S1.** Mapping mutations associated with rescued fertility in the *EMS255* line by whole-genome sequencing. The charts show the positions of *de novo* mutations on individual *Arabidopsis* chromosomes (x-axis) and their frequency in the B2 population exhibiting increased fertility (y-axis, %). As the causative mutation is recessive, it is expected to be fully associated with the rescued phenotype. The orange circle on chromosome 4 indicates the mutation in the At4g17270 locus.

```
HsMST3    36   FTKLEKIGKGSFGEVFKGIDNRTQKVVAIKIIDLEEAEDEIEDIQQEITVLSQCDSPYVT  95
               F++ E IG+GSFG+V+K  D    K VAIK+IDLEE+EDEIEDIQ+EI+VLSQC  PY+T
AtMAPK4K1 15   FSQFELIGRGSFGDVYKAFDTELNKDVAIKVIDLEESEDEIEDIQKEISVLSQCRCPYIT  74

HsMST3    96   KYYGSYLKDTKLWIIMEYLGGGSALDLLEPG-PLDETQIATILREILKGLDYLHSEKKIH  154
               +YYGSYL  TKLWIIMEY+ GGS  DLL+PG PLDE  IA I R++L  ++YLH+E KIH
AtMAPK4K1 75   EYYGSYLHQTKLWIIMEYMAGGSVADLLQPGNPLDEISIACITRDLLHAVEYLHAEGKIH  134

HsMST3    155  RDIKAANVLLSEHGEVKLADFGVAGQLTDTQIKRNTFVGTPFWMAPEVIKQS-AYDSKAD  213
               RDIKAAN+LLSE+G+VK+ADFGV+ QLT T  +R TFVGTPFWMAPEVI+ S  Y+ KAD
AtMAPK4K1 135  RDIKAANILLSENGDVKVADFGVSAQLTRTISRRKTFVGTPFWMAPEVIQNSEGYNEKAD  194

HsMST3    214  IWSLGITAIELARGEPPHSELHPMKVLFLIPKNNPPTLEGNYSKPLKEFVEACLNKEPSF  273
               IWSLGIT IE+A+GEPP ++LHPM+VLF+IP+ +PP L+ ++S+PLKEFV  CL K P+
AtMAPK4K1 195  IWSLGITMIEMAKGEPPLADLHPMRVLFIIPRESPPQLDEHFSRPLKEFVSFCLKKAPAE  254

HsMST3    274  RPTAKELLKHKFILRNAKKTSYLTELIDRYKRWKAEQSHDDSSSEDSDAETDGQASGGSD  333
               RP AKELLKH+FI +NA+K+  L E I    +++ +        ED +  T+G  +
AtMAPK4K1 255  RPNAKELLKHRFI-KNARKSPKLLERIRERPKYQVK--------EDEEIPTNGPKAPAES  305

HsMST3    334  SG  335
               SG
AtMAPK4K1 306  SG  307
```

**Figure S2.** The MST3-MO25β interaction interphase is evolutionary conserved. Protein sequence alignment of human MST3 and *Arabidopsis thaliana* MAPK4K1. Amino acid residues comprising the MST3-MO25β interaction interphase, as determined by crystallography (Mehellou et al., 2013), are highlighted in red.

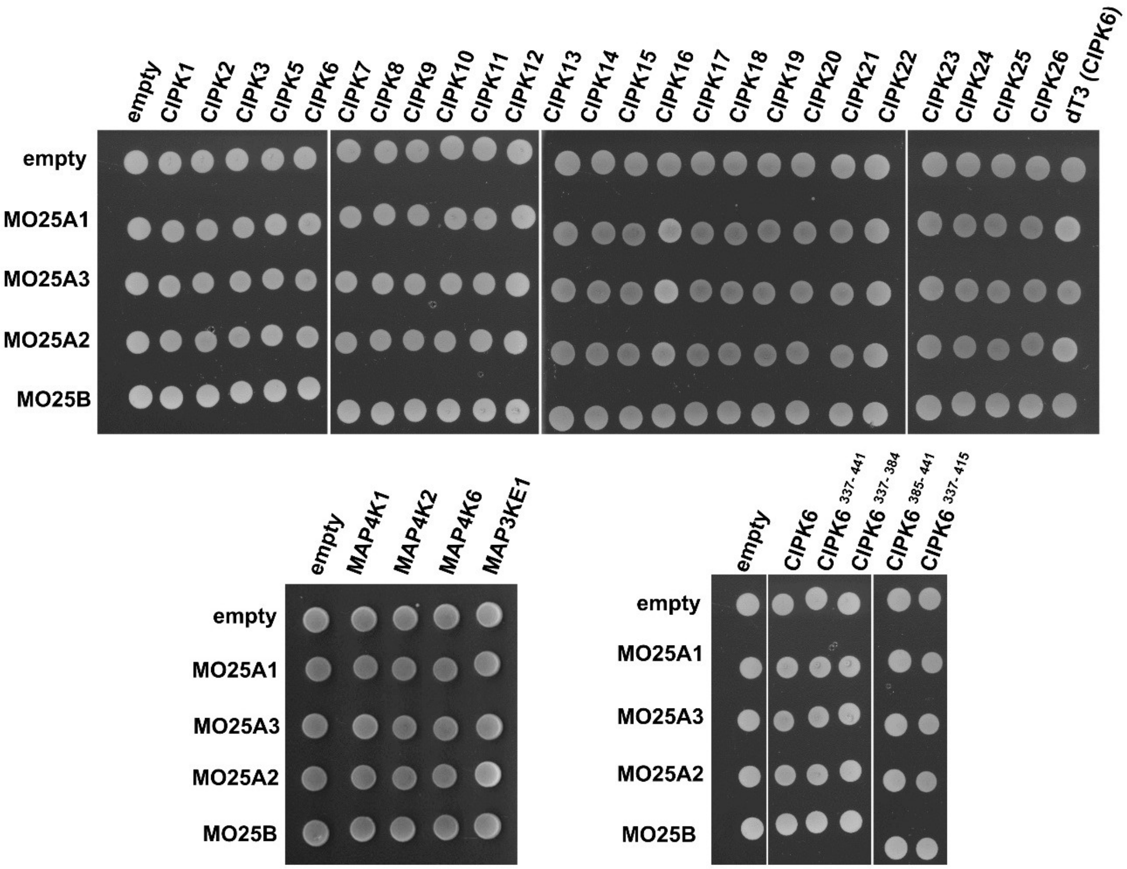


**Figure S3**. Mating controls for the yeast two-hybrid assays shown in Figure 6. Yeast colonies were grown on SD-LT plates.

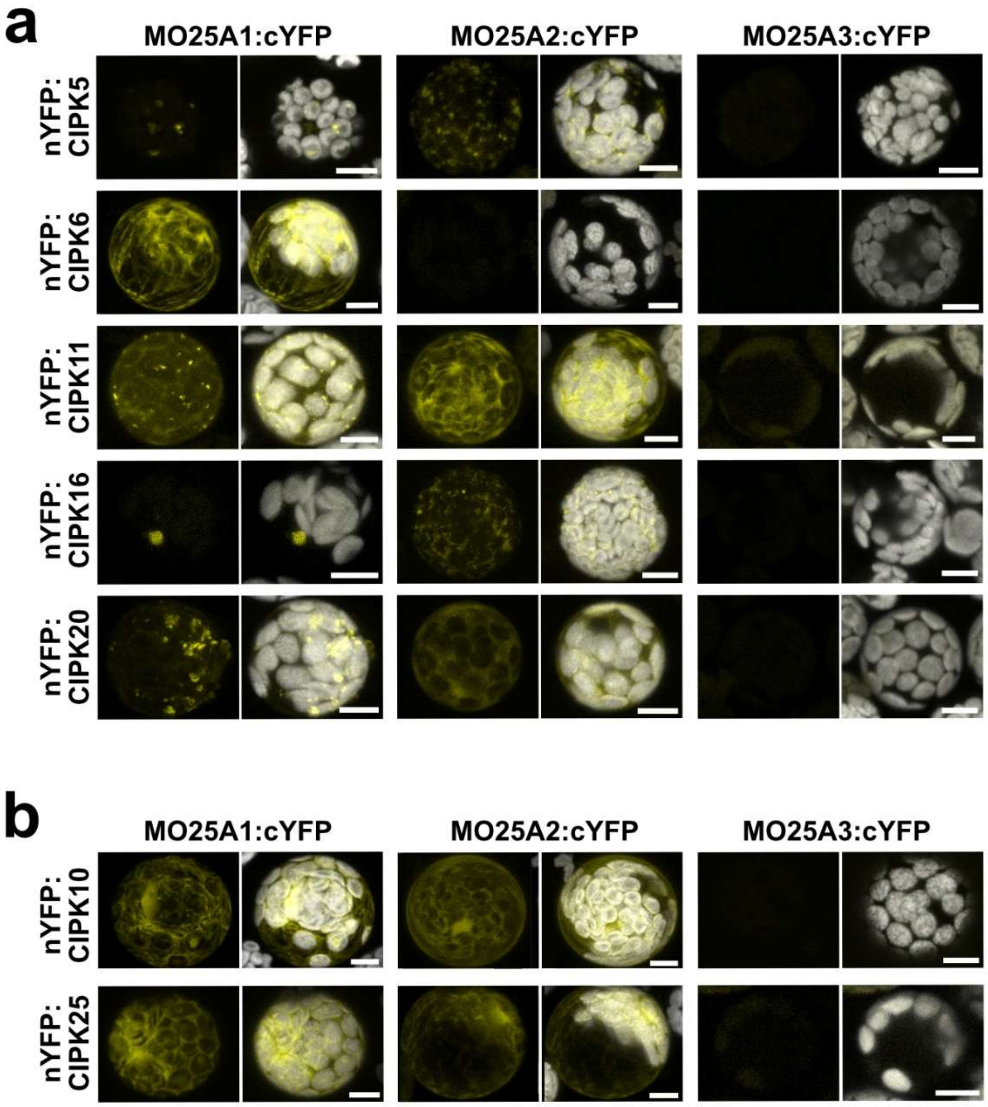


**Figure S4**. Bimolecular fluorescence complementation assay between MO25A proteins and CIPK kinases identified by yeast two-hybrid assay (a) or selected based on a conserved C-terminal sequence motif (b). Scale bars = 5 µm.

| | | |
|---|---|---|
| interacting in Y2H assay | CIPK5 | TLEYDRLYEEEVRPALNDIVWSWHGDNNN |
| | CIPK6 | TLEYNNFCSTALRPALKDIFWTSTPA |
| | CIPK11 | GDCYKEMWNGKLRVQLIRVCDQTSSTNAA |
| | CIPK16 | TLEYYLFCEDDVRPALKDIVWSWQGDDDE |
| | CIPK20 | TMEYKQFCDRELRPSLKDIVWKWQGNNNN |
| | CIPK1 | GDSCLYRQLYERLLKDVGTSSPE |
| | CIPK2 | TMEYQKLVEEDLRPALADIVWVWQGEKEK |
| | CIPK3 | EGEEEEEEVESHWSELIVELEEIVLSWHN |
| | CIPK7 | VKVVEGEEDDSRWSDLITELEDIVLSWHNDIM |
| | CIPK8 | GDAEEYLKFYKTFCSKLDDIIWKPPDASMR |
| | CIPK9 | DTLEFHKFYKNFSSGLKDVVWNTDAAAEE |
| | CIPK10 | TLEYQKLVAEDLRPALSDIVWVWQGEKDE |
| | CIPK12 | KTEYEDFCNNELKPKLQNLTADDVVAEPV |
| | CIPK13 | IEEYEEFCNKELRPQLEKLMHYQADEVEE |
| | CIPK14 | AASGRDLWTDTLRPFFVELVHESDQTDPE |
| | CIPK15 | TAEYKRVMKESIRPALIDFVLAWH |
| | CIPK17 | GDSSLYKQLYERLLNELGSSSQV |
| | CIPK18 | REEYEEFCNKELRPELEKLIHEEVVV |
| | CIPK19 | KMEYDEFCNKELKPKLQNLSSENGQRVSG |
| | CIPK21 | VIEISKSAGELRLYMEFCQSLSSLLTAEVS |
| | CIPK22 | QRDVGLVWADALRQKLRRLINQPVYKVPD |
| | CIPK23 | TLEFHKFY-KNLTTGLKDIVWKTIDEEKE |
| | CIPK24 | TLEYHKFYKKLCS-KLENIIWRATEGIPK |
| | CIPK25 | TLEYNRLYEEHVRPALEEIVWSWHGDNHN |
| | CIPK26 | TLEFHKFYKKLSTS-LNDVVWKSGESSGL |

**Figure S5.** Sequence alignment of the C-terminal regions of Arabidopsis CIPKs. Residues conserved among MO25A-interacting CIPKs are highlighted in red.

| AGI code | Gene name | Annotation | oligo dT library | Random primer library |
|---|---|---|---|---|
| AT4G30960 | CIPK6 | CBL-interacting protein kinase 6 | 10 (3) | 9 (2) |
| AT5G10930 | CIPK5 | CBL-interacting protein kinase 5 | 1 | |
| AT5G45820 | CIPK20 | CBL-interacting protein kinase 20 | 4 (1) | |
| AT1G51200 | SAP2 | A20/AN1-like zinc finger family protein | | 1 |
| AT3G03790 | RCC1 | ankyrin repeat family protein / regulator of chromosome condensation family protein | | 2(1) |

**Table S1.** Proteins identified in yeast two-hybrid screens using MO25A1 as bait against two cDNA libraries. The table lists the number of clones identified for each gene, with the corresponding number of unique clones shown in parentheses.
